# Myogenetic oligodeoxynucleotides as anti-nucleolin aptamers inhibit the growth of embryonal rhabdomyosarcoma cells

**DOI:** 10.1101/2021.10.18.464889

**Authors:** Naoki Nohira, Sayaka Shinji, Shunichi Nakamura, Yuma Nihashi, Takeshi Shimosato, Tomohide Takaya

**Affiliations:** Department of Agricultural and Life Sciences, Faculty of Agriculture, Shinshu University, Nagano, Japan; Department of Agriculture, Graduate School of Science and Technology, Shinshu University, Nagano, Japan; Department of Science and Technology, Graduate School of Medicine, Science and Technology, Shinshu University, Nagano, Japan; Department of Biomolecular Innovation, Institute for Biomedical Sciences, Shinshu University, Nagano, Japan

**Author notes:** Nohira Naoki, Sayaka Shinji, and Shunichi Nakamura equally contributed to this work. Correspondence, Department of Agricultural and Life Sciences, Faculty of Agriculture, Shinshu University, 8304 Minami-minowa, Kami-ina, Nagano 399-4598, Japan.

**Keywords:** Aptamer, Embryonal rhabdomyosarcoma, Myogenetic oligodeoxynucleotide, Nucleolin

## Abstract

**Background:** Embryonal rhabdomyosarcoma (ERMS) is the muscle-derived tumor retaining myogenic ability. iSN04 and AS1411, which are myogenetic oligodeoxynucleotides (myoDNs) serving as anti-nucleolin aptamers, have been reported to inhibit the proliferation and induce the differentiation of myoblasts. The present study investigated the effects of iSN04 and AS1411 on the growth of multiple ERMS1 cell lines in vitro.

**Methods:** Three patient-derived ERMS cell lines, ERMS1, KYM1, and RD, were used. Nucleolin expression and localization in these cells was confirmed by immunostaining. The effects of iSN04 or AS1411 on the growth of ERMS cells were examined by cell counting, EdU staining, quantitative RT-PCR (qPCR), and three-dimensional culture of tumorspheres.

**Results:** In all ERMS cell lines, nucleolin was abundantly expressed, and localized and concentrated in nucleoli, similar to myoblasts. Both iSN04 and AS1411 (10-30 μM) significantly decreased the number of all ERMS cells; however, their optimal conditions were different among the cell lines. iSN04 (10 μM) markedly reduced the ratio of EdU^+^ cells, indicating the inhibition of cell proliferation. qPCR demonstrated that iSN04 suppressed the cell cycle, partially promoted myogenesis, but did not induce apoptosis. Finally, both iSN04 and AS1411 (10-30 μM) disrupted the formation and outgrowth of RD tumorspheres mimicking in vivo tumorigenesis.

**Conclusions:** ERMS cells expressed nucleolin, and their growth was inhibited by the anti-nucleolin aptamers, iSN04 and AS1411. The present study provides the first evidence that anti-nucleolin aptamers can be used as nucleic acid drugs for chemotherapy against ERMS.

## Background

Rhabdomyosarcoma (RMS), occurring in striated muscles throughout the body, is the most frequent soft tissue tumor in children [1]. The five-year survival rate of high-risk patients is still <30% [2], which has not been improved by standard anti-tumor chemotherapy using actinomycin D, cyclophosphamide, ifosfamide, and vincristine [1, 3]. Embryonal RMS (ERMS) is a subtype that accounts 70% of childhood RMS. A number of causative mutations, including Ras [4], Rb1 [5], and p53 [6], have been identified in ERMS. These mutations in muscular lineages, such as mesenchymal stem cells, satellite cells, or myoblasts, cause ERMS to present impaired myogenic differentiation and hyper-activated proliferation [7–9]. Therefore, it is anticipated that the agents modulating the disrupted myogenic program could be novel effective drugs for ERMS therapy.

Nucleic acid aptamers are single-strand short nucleotides that specifically bind to target molecules in a structure-dependent manner and are promising candidates for next-generation drugs against manifold diseases. Dozens of aptamers targeting tumoral proteins have been developed and their therapeutic effects on cancer cells have been studied [10]. However, there have been no reports on the treatment of RMS cells with aptamers. We recently identified a series of 18-base myogenetic oligodeoxynucleotides (myoDNs) that facilitate the differentiation of myoblasts, while suppressing the cell growth [11–13]. One of the myoDNs, iSN04, serves as the aptamer directly binding to nucleolin, improves the p53 protein level translationally inhibited by nucleolin, and finally leads to myogenesis via activation of the p53 signaling pathway [11]. Nucleolin is a ubiquitous multifunctional phosphoprotein localized in the nucleus, cytoplasm, and plasma membrane, depending on the context of cellular processes [14]. In cancers, nucleolin is frequently observed on the cell surface, which interacts with the ligands involved in proliferation and apoptosis [15]. For instance, the interaction of nucleolin with ErbB1 and Ras promotes proliferation [16], while that with Fas inhibits apoptosis [17]. Thus, several nucleolin inhibitors have been developed for application in cancer therapy. AS1411 is a 26-base guanine-rich anti-nucleolin aptamer that functions in the nucleus, cytoplasm, and plasma membrane. AS1411 has shown anti-cancer activity against acute myeloid leukemia in clinical trials [15, 18]. Intriguingly, in addition to its anti-tumor effect, AS1411 promotes myoblast differentiation to the same extent as iSN04 [11]. These findings suggest that the combined myogenetic and anti-carcinogenic abilities of anti-nucleolin aptamers might be beneficial for ERMS therapy. The present study investigated whether iSN04 and AS1411 affect the growth and myogenesis of ERMS cells.

## Methods

### Oligodeoxynucleotides

Phosphorothioated iSN04 (5’-AGA TTA GGG TGA GGG TGA-3’) was synthesized and purified by HPLC (GeneDesign, Osaka, Japan) [11–13]. AS1411 (5’-GGT GGT GGT GGT TGT GGT GGT GGT GG-3’), having a phosphodiester backbone, was synthesized and desalted (Integrated DNA Technologies, IA, USA) [11, 19]. iSN04 and AS1411 were dissolved in endotoxin-free water. An equal volume of endotoxin-free water, without iSN04 or AS1411, served as the negative control.

### Cell culture

Three ERMS cell line stocks were provided by JCRB Cell Bank (National Institutes of Biomedical Innovation, Health and Nutrition, Osaka, Japan): ERMS1 cells (JCRB1648) derived from the anaplastic pelvic ERMS of a 5-year-old female [6], KYM1 cells (JCRB0627) derived from the neck ERMS of a 9-month-old infant [20], and RD cells (JCRB9072) derived from the malignant pelvic ERMS of a 7-year-old female [21]. The ERMS cells were maintained in RPMI1640 (Nacalai, Osaka, Japan) with 10% fetal bovine serum (HyClone; GE Healthcare, UT, USA), 100 units/ml penicillin, and 100 μg/ml streptomycin at 37°C with 5% CO_2_ [22]. The commercially available human myoblasts isolated from the healthy subject (CC-2580; Lonza, MD, USA) were maintained in Skeletal Muscle Growth Media-2 (CC-3245; Lonza) on the dishes coated with collagen type I-C (Cellmatrix; Nitta Gelatin, Osaka, Japan) at 37°C with 5% CO_2_ [11, 12, 23].

### Immunocytochemistry

The cells were fixed with 2% paraformaldehyde, permeabilized with 0.2% Triton X-100, and immunostained with 1.0 μg/ml rabbit polyclonal anti-nucleolin antibody (ab22758; Abcam, Cambridge, UK). 0.1 μg/ml Alexa Fluor 488-conjugated donkey polyclonal anti-rabbit IgG antibody (Jackson ImmunoResearch, PA, USA) was used as a secondary antibody. Cell nuclei were stained with DAPI (Nacalai). Fluorescent images were captured using an EVOS FL Auto microscope (AMAFD1000; Thermo Fisher Scientific, MA, USA).

### Cell counting

5.0×10^4^ cells/well were seeded on 12-well (ERMS1 and RD) or 24-well (KYM1) plates and then treated with iSN04 or AS1411 the next day. After 72 h (ERMS1 and KYM1) or 92 h (RD) of treatment, when the control group became confluent, the cells were completely dissociated by 0.25% trypsin and subjected to cell counting using a hemocytometer.

### EdU staining

1.0×10^5^ cells (ERMS1 and RD) or 3.0×10^5^ cells (KYM1) were seeded on 30-mm dishes and treated with 10 μM iSN04 the next day. After 24 h (ERMS1 and KYM1) or 48 h (RD) of the treatment, EdU (5-ethynyl-2’-deoxyuridine) was added at a final concentration of 10 μM, and the cells were cultured for 3 h. EdU staining was performed using the Click-iT EdU Imaging Kit (Thermo Fisher Scientific), according to the manufacturer’s instruction. Cell nuclei were visualized by DAPI staining. The ratio of EdU^+^ cells was defined as the number of EdU^+^nuclei divided by the total number of nuclei using ImageJ software (National Institutes of Health, USA) [23].

### Quantitative real-time RT-PCR (qPCR)

1.5×10^5^ cells (RD), 2.0×10^5^ cells (ERMS1), or 3.0×10^5^ cells (KYM1) were seeded on 30-mm dishes and treated with 10 μM iSN04 the next day. After 72 h (ERMS1 and KYM1) or 96 h (RD) of treatment, the total RNA from the cells was isolated using NucleoSpin RNA Plus (Macherey-Nagel, Düren, Germany) and reverse transcribed using ReverTra Ace qPCR RT Master Mix (TOYOBO, Osaka, Japan). qPCR was performed using GoTaq qPCR Master Mix (Promega, WI, USA) with the StepOne Real-Time PCR System (Thermo Fisher Scientific). The amount of each transcript was normalized to that of *GAPDH*. The results are presented as fold-changes. The primer sequences are listed in Table 1.

**Table 1.**
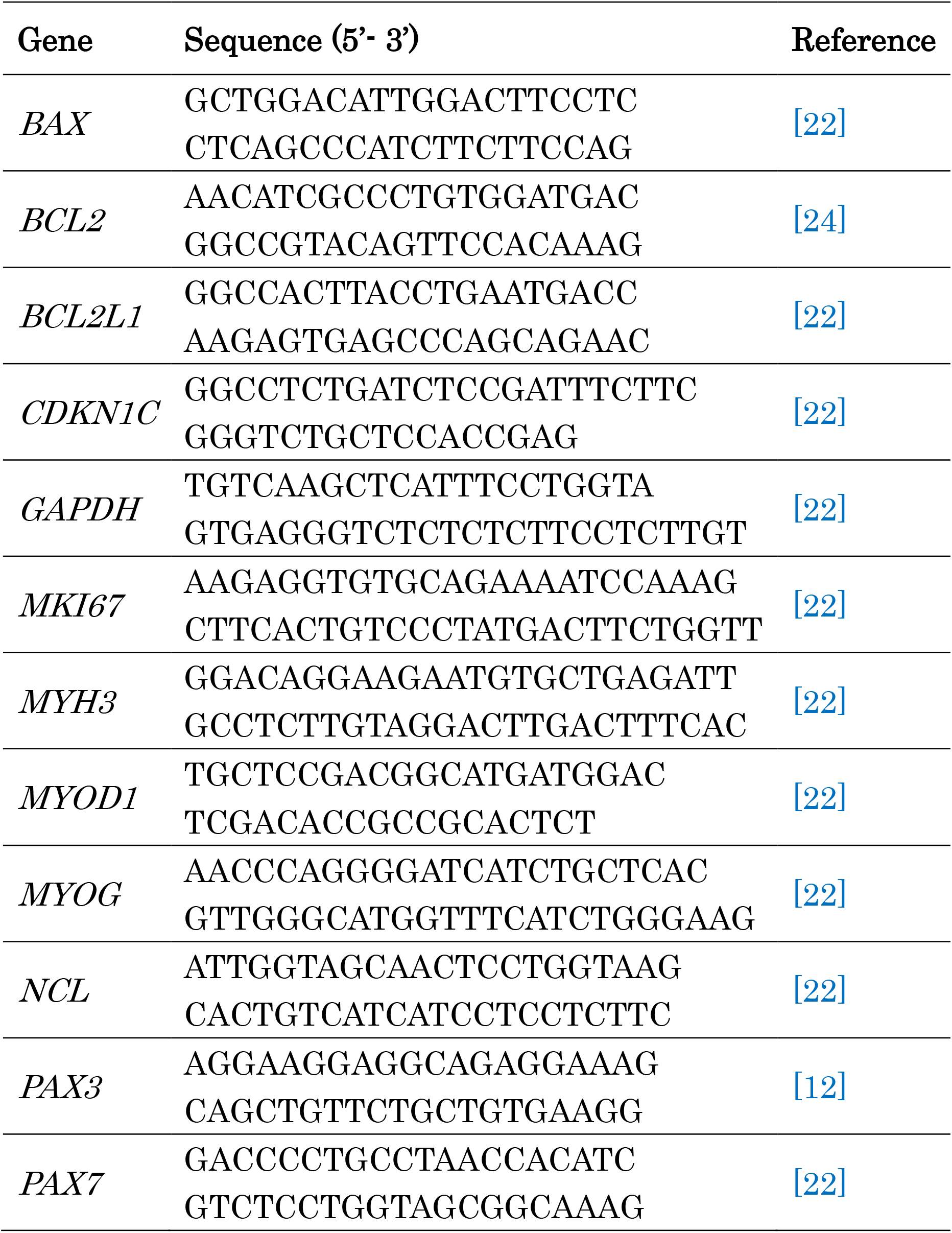
Primer sequences for qPCR

### Three-dimensional (3D) culture of RD tumorspheres

RD cells were dissociated and suspended in 3D Tumorsphere Medium XF (PromoCell, Heidelberg, Germany). Drops (300 cells/30 μl) were placed on 24-well floating-culture plates (Sumitomo Bakelite, Tokyo, Japan).

Subsequently, the plates were turned over for the hanging-drop culture. After 3 days, the plates were turned over again, and 300 μl/well of 3D Tumorsphere Medium XF with 10 or 30 μM of iSN04 or AS1411 was added to RD tumorspheres (defined as day 0). The spheres were maintained without medium exchange for 10 days. Bright-field images were taken using an EVOS FL Auto microscope [22].

### Statistical analysis

Results are presented as the mean ± standard error. Statistical comparisons were performed using unpaired two-tailed Student’s *t*-test or multiple comparison test with Scheffe’s *F* test following one-way analysis of variance. Statistical significance was set at a *p* value < 0.05.

## Results

### Nucleolin expression and localization in ERMS cells

Three patient-derived ERMS cell lines, ERMS1 [6], KYM1 [20], and RD [21], were used in this study. These cells are morphologically different from each other and from myoblasts, reflecting the diverse phenotypes of ERMS cells (Fig. 1A). Initially, the expression and localization of nucleolin in these cells was confirmed because there has not yet been any research on nucleolin in RMS cells. RT-PCR indicated that nucleolin (*NCL*) mRNA was abundantly transcribed in all ERMS cell lines as well as myoblasts (Fig. 1B). Immunostaining revealed that nucleolin localized intensively in the nucleoli, but not in the cytoplasm and on the plasma membrane of all ERMS cells (Fig. 1C). The localization patterns of nucleolin in these cells were similar to those observed in myoblasts.

**Fig. 1.**
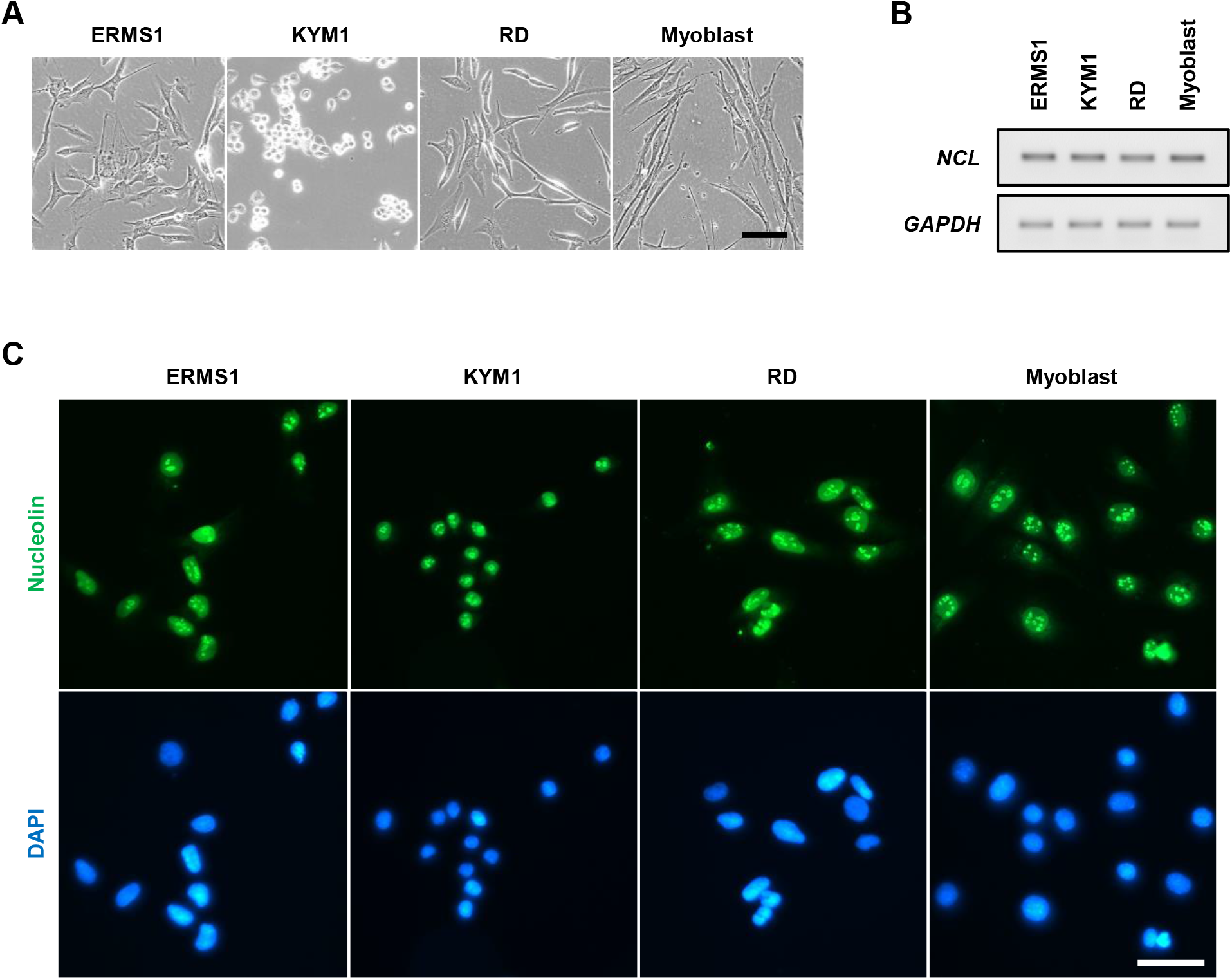
Expression and localization of nucleolin in ERMS cells. (**A**) Representative phase-contrast images of ERMS cells and myoblasts. Scale bar, 100 μm. (**B**) RT-PCR result of nucleolin (*NCL*) expression in ERMS cells and myoblasts. (**C**) Representative immunofluorescent images of nucleolin staining of ERMS cells and myoblasts. Scale bar, 50 μm.

### iSN04 and AS1411 inhibit the growth of ERMS cells

Next, the effects of anti-nucleolin aptamers on the growth of ERMS cells were investigated. The cells were treated with iSN04 or AS1411 until their negative controls became confluent, and the number of cells was counted (Fig. 2A). Both iSN04 and AS1411 significantly decreased the number of all ERMS cells compared to the control group. In ERMS1 cells, the inhibitory effects of both iSN04 and AS1411 were saturated at a concentration of 10 μM; however, the activity of AS1411 was markedly higher than that of iSN04. In contrast, KYM1 cells were more sensitive to iSN04 than to AS1411. In RD cells, 10 and 30 μM of iSN04 and AS1411 significantly reduced the cell numbers in a dose-dependent manner, and AS1411 showed higher activity compared to iSN04. These results demonstrate that anti-nucleolin aptamers are effective in inhibiting the growth of ERMS cells, but their activities or sensitivities to the cells may differ among cell lines.

**Fig. 2.**
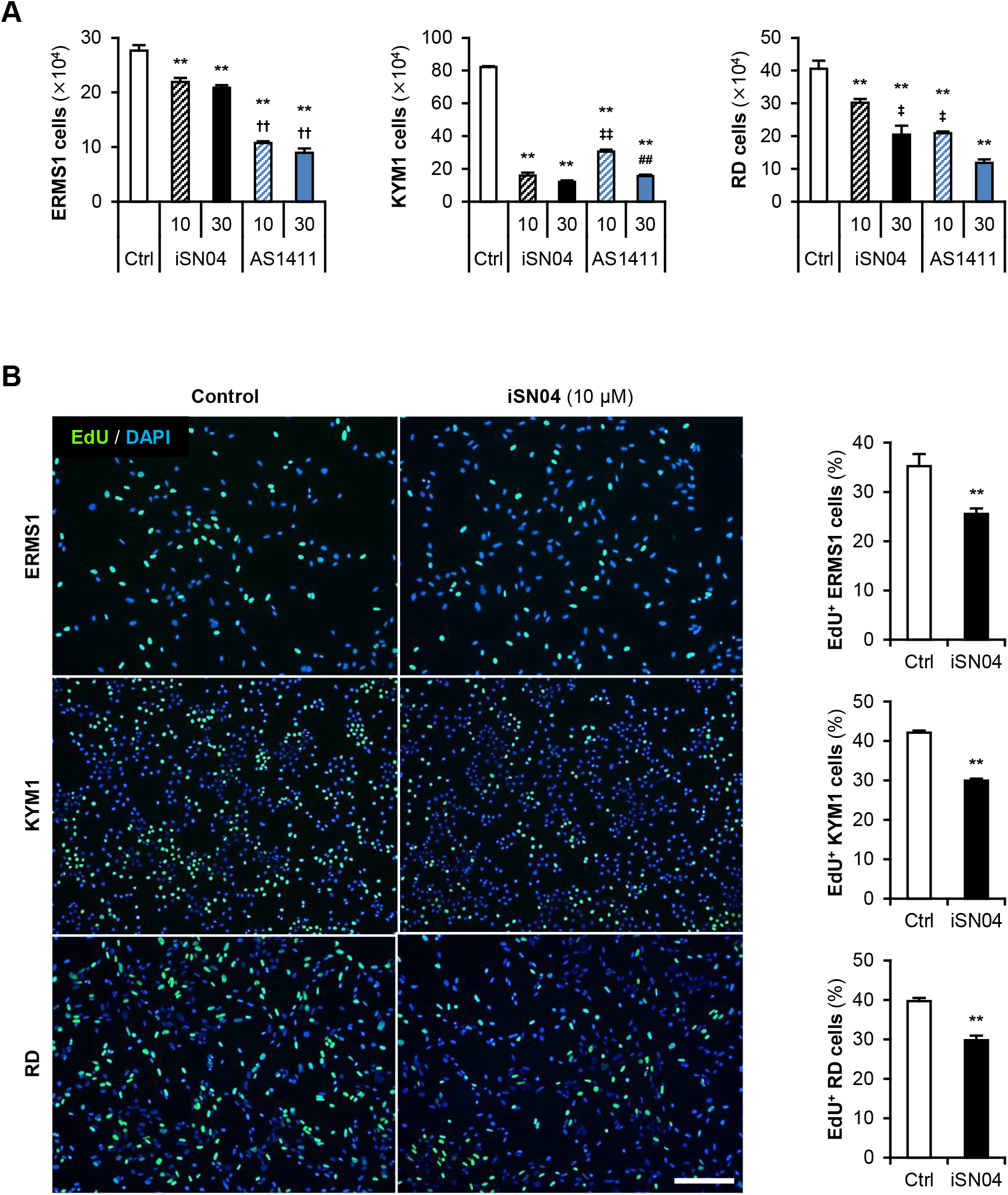
Nucleolin aptamers inhibit the growth of ERMS cells. (**A**) Numbers of the ERMS cells treated with 10 or 30 μM of iSN04 or AS1411 for 72 h (ERMS1 and KYM1) or 96 h (RD). ^**^ *p* < 0.01 vs control; ^††^ *p* < 0.01 vs 30 μM iSN04; ^‡^ *p* < 0.05, ^‡‡^ *p* < 0.01 vs 10 μM iSN04; ^##^ *p* < 0.01 vs 10 μM AS1411 (Scheffe’s *F* test). *n* = 4. (**B**) Representative images of the ERMS cells treated with 10 μM iSN04 for 24 h (ERMS1 and KYM1) or 48 h (RD). Scale bar, 200 μm. Ratio of EdU^+^ cells were quantified. ^**^ *p* < 0.01 vs control (Student’s *t*-test). *n* = 6.

To clarify the action of anti-nucleolin aptamers, ERMS cells treated with iSN04 were subjected to EdU staining, and the EdU^+^ cells replicating genomic DNA were quantified (Fig. 2B). In all ERMS cell lines, iSN04 markedly decreased the ratio of EdU^+^ cells, indicating that iSN04 delayed the cell cycle. These results demonstrate that iSN04 suppresses cell proliferation, resulting in inhibitory effects on the growth of ERMS cell lines.

### iSN04 alters the gene expression in ERMS cells

The effects of iSN04 on the gene expression in ERMS cells were quantified by qPCR (Fig. 3). iSN04 significantly increased the mRNA levels of the cyclin-dependent kinase inhibitor 1C (p57^Kip2^) (*CDKN1C*) in all ERMS cell lines, and decreased the transcription of proliferation marker Ki-67 (*MKI67*) in ERMS1 and RD cells, which corresponded well with the results of EdU staining. The mRNA levels of apoptosis-related factors, Bax (*BAX*), Bcl-2 (*BCL2*), and Bcl-xL (*BCL2L1*), were not altered by iSN04 in every ERMS cell line, except for *BCL2L1* in ERMS1 cells. These data demonstrate that iSN04 attenuates the growth of ERMS cells by inhibiting the cell cycle, but not by inducing apoptosis.

**Fig. 3.**
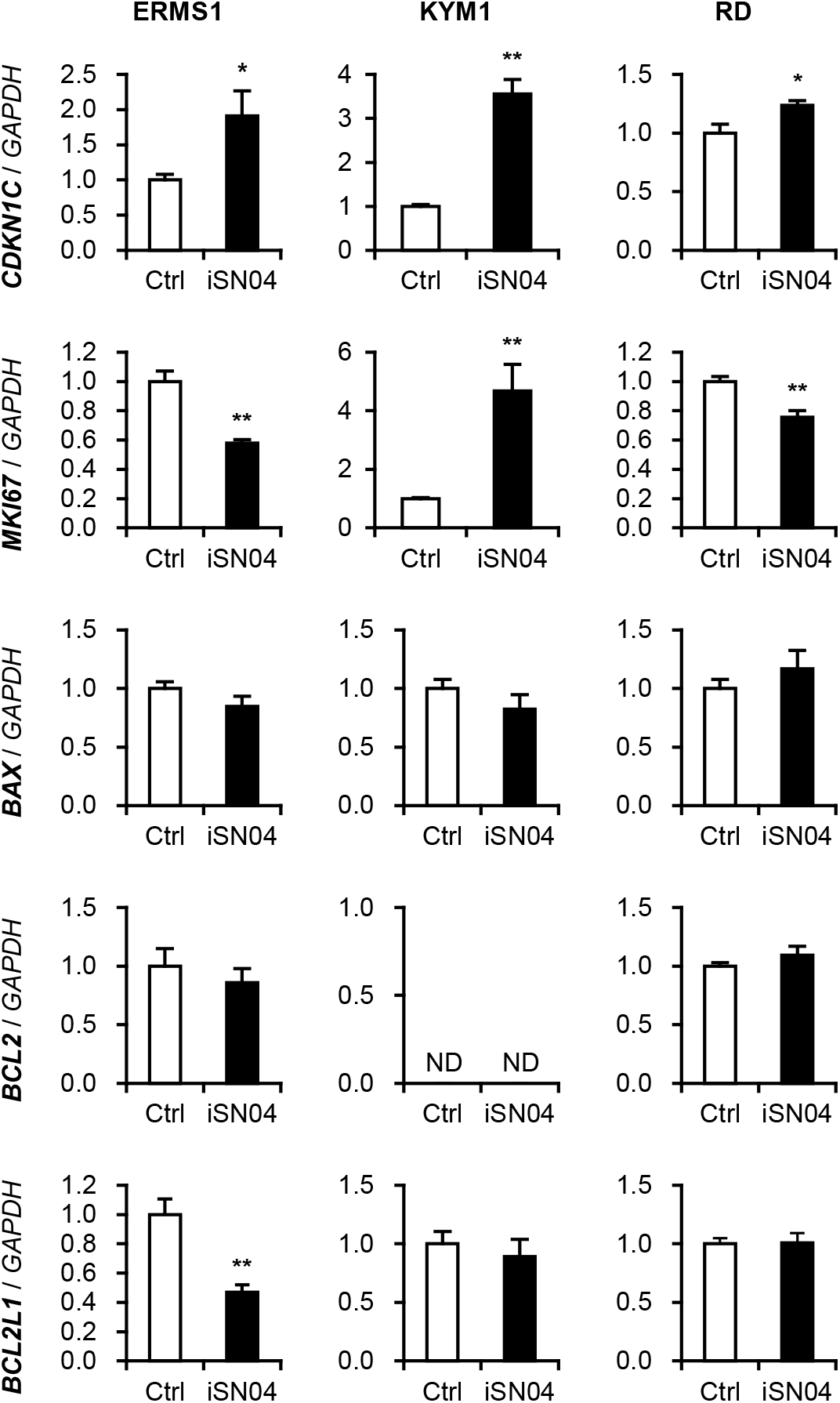
Effects of iSN04 on the proliferative and apoptotic gene expression in ERMS cells. qPCR quantified the mRNA levels of p57^Kip2^ (*CDKN1C*), Ki-67 (*MKI67*), Bax (*BAX*), Bcl-2 (*BCL2*), and Bcl-xL (*BCL2L1*) in the ERMS cells treated with 10 μM iSN04 for 72 h (ERMS1 and KYM1) or 96 h (RD). Mean value of control group was set to 1.0 in each gene. ND, not detected. ^*^ *p* < 0.05, ^**^ *p* < 0.01 vs control (Student’s *t*-test) *n* = 4.

iSN04, as a myoDN, not only suppresses the proliferation, but also promotes the myogenic differentiation of myoblasts [11–13]. Since ERMS cells retain myogenic abilities even though they are dysregulated [8], we investigated whether iSN04 upregulates the myogenic gene expression in ERMS cells (Fig. 4). In RD cells, iSN04 significantly decreased the mRNA levels of Pax3 (*PAX3*) and Pax7 (*PAX7*), which are undifferentiated myogenic transcription factors. Although iSN04 did not change the expression of MyoD (*MYOD1*) as a master regulator of myogenic program, it markedly induced myogenin (*MYOG*) as a myogenic transcription factor in KYM1 cells and embryonic myosin heavy chain (MHC) (*MYH3*) as a sarcomeric protein in RD cells. Overall, iSN04 tended to induce myogenic differentiation, but its effects were divergent among ERMS cell lines.

**Fig. 4.**
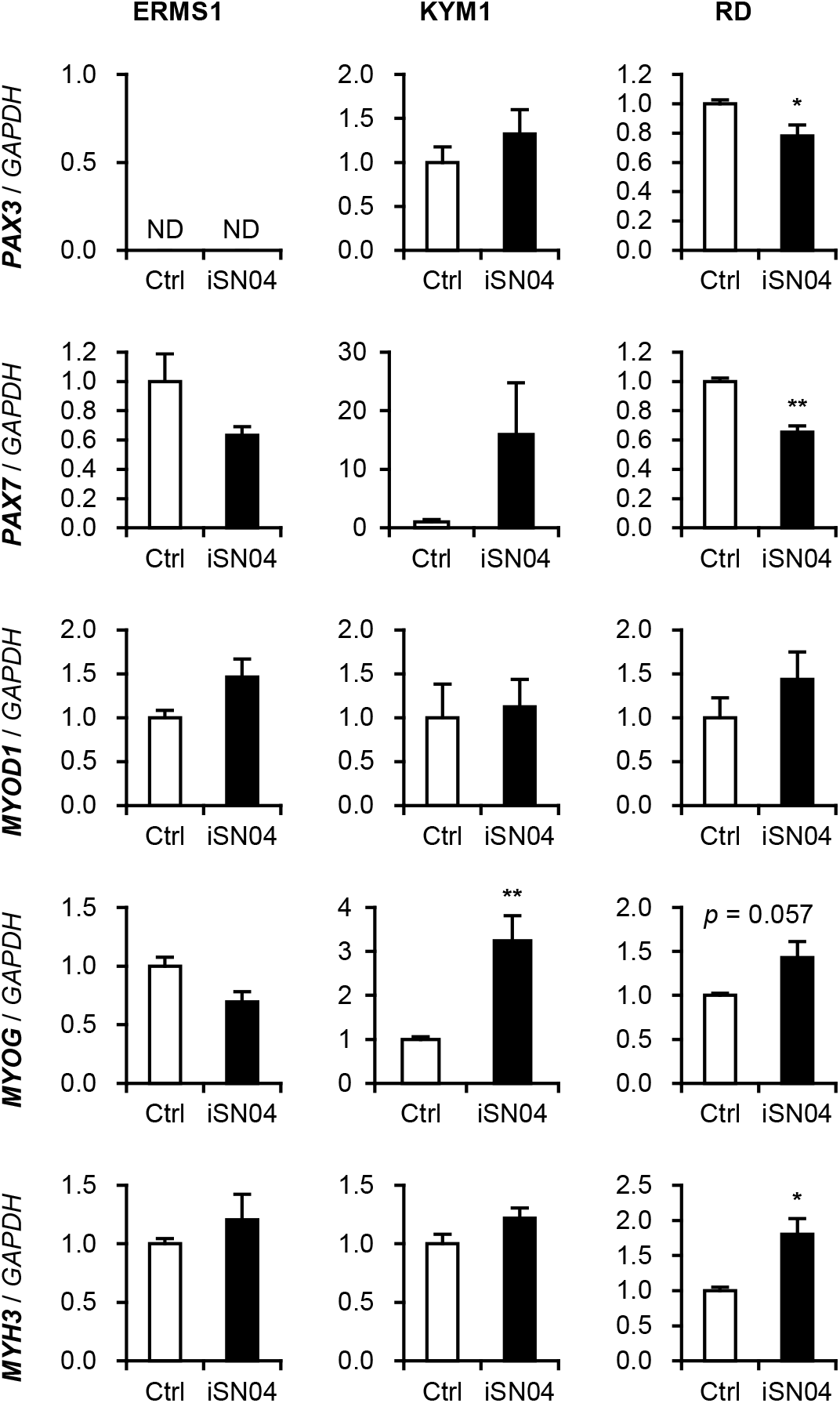
Effects of iSN04 on the myogenic gene expression in ERMS cells. qPCR quantified the mRNA levels of Pax3 (*PAX3*), Pax7 (*PAX7*), MyoD (*MYOD1*), myogenin (*MYOG*), and embryonic MHC (*MYH3*) in the ERMS cells treated with 10 μM iSN04 for 72 h (ERMS1 and KYM1) or 96 h (RD). Mean value of control group was set to 1.0 in each gene. ND, not detected. ^*^ *p* < 0.05, ^**^*p* < 0.01 vs control (Student’s *t*-test) *n* = 4.

### iSN04 and AS1411 disturb the formation of RD tumorspheres

The 3D-culture of tumorspheres is a valuable method to mimic in vivo tumorigenesis because the cells in spheres can exert inherent characteristics of cancer stem cells [25]. We have recently developed a xeno-free floating culture system for tumorspheres of RD cells [22]. The initial RD aggregation formed by hanging-drop culture was subsequently subjected to floating culture with iSN04 or AS1411. (Fig. 5). In the control group, RD tumorspheres stably grew up for 10 days and eventually shaped approximately 0.4 mm-diameter globes. However, iSN04- or AS1411-treatment disturbed the sphere formation. AS1411 interfered with the growth of some RD spheres, which indicated abnormal shapes and small diameters. iSN04 disrupted the formation and growth of RD tumorspheres more severely than AS1411. iSN04-treatment arrested the outgrowth of RD aggregation and broke some of them in small clusters. These results suggest that anti-nucleolin aptamers are effective in suppressing ERMS tumorigenesis, particularly in small metastatic lesions.

**Fig. 5.**
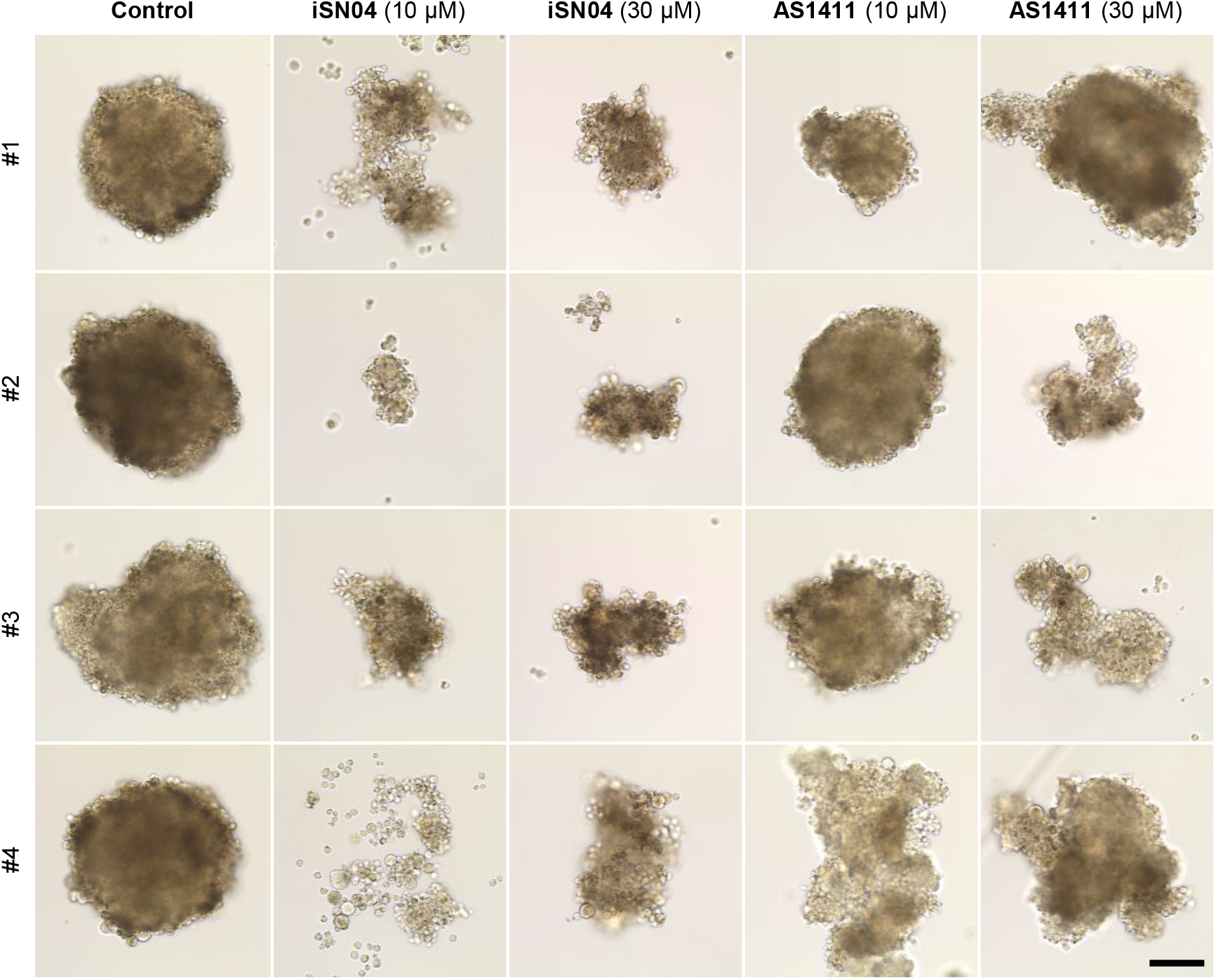
iSN04 and AS1411 disturb the formation of RD tumorspheres. Representative bright-field images of the RD tumorspheres treated with 10 or 30 μM of iSN04 or AS1411 for 10 days. Scale bar, 100 μm.

## Discussion

The present study indicated that multiple ERMS cell lines express nucleolin, and their growth is inhibited by anti-nucleolin aptamers, iSN04 and AS1411. The role of nucleolin in RMS has not yet been reported. In sarcoma, it has been only studied that nucleolin is involved in GSK3β-mediated stability of HIF1α mRNA in osteosarcoma cells [26]. The biological function of nucleolin in sarcoma needs to be further examined. Fortuitously, a number of cancer studies have provided evidence for the contribution of nucleolin in tumorigenesis. The localization and function of nucleolin varies in cancers [15]. Cell surface nucleolin is involved in ErbB1- and Ras-regulated proliferation [16], and it blocks Fas-induced apoptosis [17]. Cytoplasmic nucleolin binds to mRNAs of Bcl-2 [27] and Bcl-xL [28] to stabilize them, thereby protecting cancers against apoptosis. However, in ERMS cells, nucleolin was not detected on the plasma membrane or in the cytoplasm. Correspondingly, antagonizing nucleolin by iSN04 did not alter the mRNA levels of Bcl-2 and Bcl-xL. In ERMS cells, nucleolin was localized in the nucleoli. Nucleolar nucleolin has been reported to interact with ribosomal DNA (rDNA) and increase the RNA polymerase I transcription [29]. This process is commonly activated in tumors because hyperproliferative cancer cells require a large amount of protein synthesis. Therefore, nucleolar nucleolin is considered to play an indispensable role in the growth of cancer cells [15]. This could be one of the mechanisms by which anti-nucleolin aptamers attenuate the growth of ERMS cells. Both iSN04 and AS1411 are speculated to interact with the RNA-binding domains of nucleolin [11, 13, 18]. These aptamers might trap nucleolar nucleolin by competing with rDNA in ERMS cells.

In general, aptamers are set to target the membrane or extracellular proteins in view of drug action and are developed by the systemic evolution of ligands by exponential enrichment (SELEX) [30]. In contrast, iSN04 was identified as a myoDN that promotes myogenic differentiation [11], and AS1411 was discovered as an anti-proliferative nucleotide [18]. Nuclear nucleolin was defined as the target a posteriori. Both 18-base iSN04 and 26-base AS1411 have been confirmed to be incorporated into the cytoplasm without any carriers or transfection reagents [11, 18], probably because of their short sequences compared to the average length of the known aptamers (51 bases) [31]. Single-strand short DNA is usually taken up by cells through the endocytic process termed gymnosis [32], which enables compact DNA aptamers to target cytoplasmic and nuclear proteins. In addition to the anti-cancer aptamers working outside the cells [10], iSN04 and AS1411 present alternative directions for the development of therapeutic aptamers targeting the intracellular factors. It should be surmounted that working concentrations of iSN04 and AS1411 (10-30 μM) were relatively high compared to those of small molecules, probably because only a part of them is up taken into cytoplasm. To improve cell permeabilities of anti-nucleolin aptamers, drug delivery systems such as lipid nanoparticles need to be tested.

Intriguingly, the inhibitory activities of iSN04 and AS1411 on proliferation were different among the ERMS cell lines. This provides an important insight into the development of custom aptamers for tailor-made medicines. The previous studies suggested that AS1411 recognizes only a small subset of the total nucleolin [18, 33], because nucleolin is post-transcriptionally modified and forms a complex with various partners. Certain forms of nucleolin interplaying iSN04 and AS1411 might differ, which would impact the drug efficacy. This is also related to the myogenetic activity of iSN04 in ERMS cells. We recently demonstrated that both iSN04 and AS1411 robustly facilitate the myogenic differentiation of myoblasts [11–13]. However, induction of myogenic gene expression by iSN04 was partial and varied among the ERMS cell lines. In addition to the dysregulated myogenic program in ERMS cells, the presence of nucleolin might be distinct from that in myoblasts. To overcome these issues, it is necessary to enhance the myogenetic activity of anti-nucleolin aptamers in ERMS cells. For example, a histone deacetylase inhibitor (HDACI), trichostatin A, promotes myoblast differentiation by upregulating myogenic gene transcription [34]. Another HDACI, PXD-101, has been reported to induce myogenesis in RD cells by increasing MyoD, myogenin, and MHC expression [35]. Epigenetic alteration by HDACIs has the potential to synergistically improve the myogenetic activity of anti-nucleolin aptamers. The current standard chemotherapy for RMS is the combination treatment of multiple anti-tumor drugs [1, 3]. For clinical application, the co-effects of anti-nucleolin aptamers with existing agents need to be validated in the future.

## Conclusion

This is the first study to report the expression and inhibition of nucleolin in ERMS cells. Nucleolin is localized in the nucleoli of multiple ERMS cell lines as well as in myoblasts. Two anti-nucleolin aptamers, iSN04 and AS1411, inhibited the growth of ERMS cells. Antagonizing nucleolin by iSN04 impaired the cell cycle, but did not induce apoptosis in ERMS cells. iSN04 partially promoted myogenic transformation by regulating the myogenic gene expression. These results indicate that anti-nucleolin aptamers can be used as alternatives or potential drug candidates for chemotherapy against ERMS.

## Abbreviations

3D: Three-dimensional
ERMS: Embryonal rhabdomyosarcoma
HDACI: Histone deacetylase inhibitor
MHC: Myosin heavy chain
myoDN: Myogenetic oligodeoxynucleotide
RMS: rhabdomyosarcoma

## Acknowledgments

The preprint has been posted on bioRxiv.

## Authors’ contributions

T.T. designed the study and wrote the manuscript. N.N., S.S., S.N., and Y.N. performed experiments and collected data. T.S. designed and prepared iSN04. All authors read and approved the final manuscript.

## Availability of data and materials

The datasets used during the current study are available from the corresponding author on reasonable request.

## Declarations

### Ethics approval and consent to participate

Not applicable. This study is an observational study using cell lines. The consent to participate is not applicable for the study.

### Consent for publication

Not applicable.

## References

1. Sun X, Guo W, Shen JK, Mankin HJ, Hornicek FJ, Duan Z. Rhabdomyosarcoma: Advances in molecular and cellular biology. Sarcoma, 2015; 2015: 232010. https://doi.org/10.1155/2015/232010.

2. Malempati S, Hawkins DS. Rhabdomyosarcoma: review of the Children’s Oncology Group (COG) Soft-Tissue Sarcoma Committee experience and rationale for current COG studies. Pediatr Blood Cancer, 2012; 59: 5–10. https://doi.org/10.1002/pbc.24118.

3. Chen C, Garcia HD, Scheer M, Henssen AG. Current and future treatment strategies for rhabdomyosarcoma. Front Oncol, 2019; 9: 1458. https://doi.org/10.3389/fonc.2019.01458.

4. Langenau DM, Keefe MD, Storer NY, Guyon JR, Kutok JL L. X, et al. Effects of RAS on the genesis of embryonal rhabdomyosarcoma. Genes Dev. 2007; 21: 1382–95. https://doi.org/10.1101/gad.1545007.

5. Kohashi K, Oda Y, Yamamoto H, Tamiya S, Takahira T, Takahashi Y, et al. Alterations of RB1 gene in embryonal and alveolar rhabdomyosarcoma: special reference to utility of pRB immunoreactivity in differential diagnosis of rhabdomyosarcoma subtype. J Cancer Res Clin Oncol, 2008; 134: 1097–103. https://doi.org/10.1007/s00432-008-0385-3.

6. Nishimura R, Takita J, Sato-Otsubo A, Kato M, Koh K, Hanada R, et al. Characterization of genetic lesions in rhabdomyosarcoma using a high-density single nucleotide polymorphism array. Cancer Sci, 2013; 104: 856–64. https://doi.org/10.1111/cas.12173.

7. Rubin BP, Nishijo K, Chen HI, Yi X, Schuetze DP, Pal R, et al. Evidence for an unanticipated relationship between undifferentiated pleomorphic sarcoma and embryonal rhabdomyosarcoma. Cancer Cell, 2011; 19: 177–91. https://doi.org/10.1016/j.ccr.2010.12.023.

8. Keller C, Guttridge DC. Mechanisms of impaired differentiation in rhabdomyosarcoma. FEBS J, 2013; 280: 4323–34. https://doi.org/10.1111/febs.12421.

9. Storer NY, White RM, Uong A, Price E, Nielsen GP, Langenau DM, et al. Zebrafish rhabdomyosarcoma reflects the developmental stage of oncogene expression during myogenesis. Development, 2013; 140: 3040–50. https://doi.org/10.1242/dev.087858.

10. Li Z, Fu X, Huang J, Zeng P, Huang Y, Chen X, et al. Advances in screening and development of therapeutic aptamers against cancer cells. Front Cell Dev Biol, 2021; 9: 662791. https://doi.org/10.3389/fcell.2021.662791

11. Shinji S, Umezawa K, Nihashi Y, Nakamura S, Shimosato T, Takaya T. Identification of the myogenetic oligodeoxynucleotides (myoDNs) that promote differentiation of skeletal muscle myoblasts by targeting nucleolin. Front Cell Dev Biol, 2021; 8: 616706. https://doi.org/10.3389/fcell.2020.616706.

12. Nakamura S, Yonekura S, Shimosato T, Takaya T. Myogenetic oligodeoxynucleotide (myoDN) recovers the differentiation of skeletal muscle myoblasts deteriorated by diabetes mellitus. Front Physiol, 2021; 12: 679152. https://doi.org/10.3389/fphys.2021.679152.

13. Nihashi Y, Shinji S, Umezawa K, Shimosato T, Ono T, Kagami H, et al. Myogenetic oligodeoxynucleotide complexed with berberine promotes differentiation of chicken myoblasts. Anim Sci J, 2021; 92: e13597. https://doi.org/10.1111/asj.13597.

14. Jia W, Yao Z, Zhao J, Guan Q, Gao L. New perspectives of physiological and pathological functions of nucleolin (NCL). Life Sci, 2017; 186: 1–10. https://doi.org/10.1016/j.lfs.2017.07.025.

15. Berger CM, Gaume X, Bouvet P. The roles of nucleolin subcellular localization in cancer. Biochimie, 2015; 113: 78–85. https://doi.org/10.1016/j.biochi.2015.03.023.

16. Farin K, Schokoroy S, Haklai R, Cohen-Or I, Elad-Sfadia G, Reyes-Reyes ME, et al. Oncogenic synergism between ErbB1, nucleolin, and mutant Ras. Cancer Res, 2011; 71: 2140–51. https://doi.org/10.1158/0008-5472.CAN-10-2887.

17. Wise JF, Berkova Z, Mathur R, Zhu H, Braun FK, Tao RH, et al. Nucleolin inhibits Fas ligand binding and suppresses Fas-mediated apoptosis in vivo via a surface nucleolin-Fas complex. Blood, 2013; 121: 4729–39. https://doi.org/10.1182/blood-2012-12-471094.

18. Bates PJ, Laber DA, Miller DM, Thomas SD, Trent JO. Discovery and development of the G-rich oligonucleotide AS1411 as a novel treatment for cancer. Exp Mol Pathol, 2009; 86: 151–64. https://doi.org/10.1016/j.yexmp.2009.01.004.

19. Girvan AC, Teng Y, Casson LK, Thomas SD, Juliger S, Ball MW, et al. AGRO100 inhibits activation of nuclear factor-κB (NF-κB) by forming a complex with NF-κB essential modulator (NEMO) and nucleolin. Mol Cancer Ther, 2006; 5: 1790–9. https://doi.org/10.1158/1535-7163.MCT-05-0361.

20. Sekiguchi M, Shiroko Y, Suzuki T, Imada M, Miyahara M, Fujii G. Characterization of a human rhabdomyosarcoma cell strain in tissue culture. Biomed Pharmacother, 1985; 39: 372–80.

21. McAllister RM, Melnyk J, Finkelstein JZ, Adams EC Jr, Gardner MB. Cultivation in vitro of cells derived from a human rhabdomyosarcoma. Cancer, 1969; 24: 520–6.

22. Shinji S, Nakamura S, Nihashi Y, Umezawa K, Takaya T. Berberine and palmatine inhibit the growth of human rhabdomyosarcoma cells. Biosci Biotechnol Biochem, 2020; 84: 63–75. https://doi.org/10.1080/09168451.2019.1659714.

23. Nihashi Y, Umezawa K, Shinji S, Hamaguchi Y, Kobayashi H, Kono T, et al. Distinct cell proliferation, myogenic differentiation, and gene expression in skeletal muscle myoblasts of layer and broiler chickens. Sci Rep, 2019; 9: 16527. https://doi.org/10.1038/s41598-019-52946-4.

24. Yosef R, Pilpel N, Tokarsky-Amiel R, Biran A, Ovadya Y, Cohen S, et al. Directed elimination of senescent cells by inhibition of BCL-W and BCL-XL. Nat Commun, 2016; 7: 11190. https://doi.org/10.1038/ncomms11190.

25. He J, Xiong L, Li Q, Lin L, Miao X, Yan S, et al. 3D modeling of cancer stem cell niche. Oncotarget, 2017; 9: 1326–45. https://doi.org/10.18632/oncotarget.19847.

26. Cheng DD, Zhao HG, Yang YS. Hu T, Yang QC. GSK3β negatively regulates HIF1α mRNA stability via nucleolin in the MG63 osteosarcoma cell line. Biochem Biophys Res Commun, 2014; 443: 598–603. https://doi.org/10.1016/j.bbrc.2013.12.020.

27. Sengupta TK, Bandyopadhyay S, Fernandes DJ, Spicer EK. Identification of nucleolin as an AU-rich element binding protein involved in bcl-2 mRNA stabilization. J Biol Chem, 2004; 279: 10855–63. https://doi.org/10.1074/jbc.M309111200.

28. Zhang J, Tsaprailis G, Bowden GT. Nucleolin stabilizes Bcl-XL messenger RNA in response to UVA irradiation. Cancer Res, 2008; 68: 1046–54. https://doi.org/10.1158/0008-5472.CAN-07-1927.

29. Cong R, Das S, Ugrinova I, Kumar S, Mongelard F, Wong J, et al. Interaction of nucleolin with ribosomal RNA genes and its role in RNA polymerase I transcription. Nucleic Acids Res, 2012; 40: 9441–54. https://doi.org/10.1093/nar/gks720.

30. Wang T, Chen C, Larcher LM, Barrero RA, Veedu RN. Three decades of nucleic acid aptamer technologies: Lessons learned, progress and opportunities on aptamer development. Biotechnol Adv, 2019; 37: 28–50. https://doi.org/10.1016/j.biotechadv.2018.11.001.

31. Lee G, Jang GH, Kang HY, Song G. Predicting aptamer sequences that interact with target proteins using an aptamer-protein interaction classifier and a Monte Carlo tree search approach. PLoS One, 2021; 16: e0253760. https://doi.org/10.1371/journal.pone.0253760.

32. Juliano RL. Intracellular trafficking and endosomal release of oligonucleotides: What we know and what we don’t. Nucleic Acid Ther, 2018; 28: 166–77. https://doi.org/10.1089/nat.2018.0727.

33. Teng Y, Girvan AC, Casson LK, Pierce WM Jr, Qian M, Thormas SD, et al. AS1411 alters the localization of a complex containing protein arginine methyltransferase 5 and nucleolin. Cancer Res, 2007; 67: 10491–500. https://doi.org/10.1158/0008-5472.CAN-06-4206.

34. Hagiwara H, Saito F, Masaki T, Ikeda M, Nakamura-Ohkuma A, Shimizu T, et al. Histone deacetylase inhibitor trichostatin A enhances myogenesis by coordinating muscle regulatory factors and myogenic repressors. Biochem Biophys Res Commun, 2011; 414: 826–31. https://doi.org/10.1016/j.bbrc.2011.10.036.

35. Marampon F, Di Nisho V, Pietrantoni I, Petragnano F, Fasciani I, Scicchitano BM, et al. Pro-differentiating and radiosensitizing effects of inhibiting HDACs by PXD-101 (Belinostat) in in vitro and in vivo models of human rhabdomyosarcoma cell lines. Cancer Lett, 2019; 461: 90–101. https://doi.org/10.1016/j.canlet.2019.07.009.

